# Towards an understanding of spiral patterning in the *Sargassum muticum* shoot apex

**DOI:** 10.1101/135806

**Authors:** Marina Linardić, Siobhan A. Braybrook

## Abstract

In plants and parenchymatous brown algae the body arises through the activity of an apical meristem (a niche of cells or a single cell). The meristem produces lateral organs in specific patterns, referred to as phyllotaxis. In plants, two different control mechanisms have been proposed – one is position-dependent and relies on morphogen accumulation at future organ sites whereas the other is a lineage-based system which links phyllotaxis to the apical cell division pattern. Here we examine the apical patterning of the brown alga, *Sargassum muticum*, which exhibits spiral phyllotaxis (137.5° angle) and an unlinked apical cell division pattern. The *Sargassum* apex presents characteristics of a self-organising system, similar to plant meristems. We were unable to correlate the plant morphogen auxin with bud positioning in Sargassum, nor could we predict cell wall softening at new bud sites. Our data suggests that in *Sargassum muticum* there is no connection between phyllotaxis and the apical cell division pattern indicating a position-dependent patterning mechanism may be in place. The underlying mechanisms behind the phyllotactic patterning appear to be distinct from those seen in plants.

**Summary:** The brown alga *Sargassum muticum* displays spiral phyllotaxis developed from a position-dependent self-organising mechanism, different from that understood in plants.

## Introduction

When discussing spiral patterns in nature, D’Arcy Thompson (1917) wrote: ‘When the bricklayer builds a factory chimney, he lays his bricks in a certain steady, orderly way, with no thought of the spiral patterns to which this orderly sequence inevitably leads, and which spiral patterns are by no means “subjective”.’ This proposition, now 100 years old, implies that spiral patterns are an emergent property of local decision making processes; the underlying mechanisms may be various while the result remains the same. In developmental biology, fate decisions (such as where to place a brick, or new organ) often exhibit characteristics of emergent phenomenon. Such decisions are often made based on a position-dependent patterning system where the position of a cell within a tissue or organ specifies its fate and a signal (or morphogen) acts as an instructive agent (Scheres, 2001). An alternative mechanism depends on cell lineage, although this seems less prevalent in walled organisms such as plants (Scheres, 2001). When one examines the processes behind areal organ positioning in plants, phyllotaxis, two major theories emerge: in some early diverging land plants, phyllotactic patterning is attributed to patterned divisions at the meristematic apical cell; in spermatophytes (seed plants), they are attributed to a morphogen-based mechanism. The latter is position-dependent patterning and the former lineage-dependent.

Early diverging land plants, such as mosses and ferns, maintain a single apical cell which acts as a stem cell for the apex (Nägeli, 1845a, 1845b; Schüepp, 1926 via Korn, 1993). In mosses, the pattern of leaf production may be seen as lineage-dependent as it follows the apical cell patterning directly (Renzaglia et al., 2000; Harrison et al., 2009). In horsetails, the whorled arrangement of the leaves is independent of the division pattern in the apical cell (Golub, 1948). In fern apices, the division of the apical cell occurs successively from each face of a three-sided apical cell producing packets of daughter cells at 120° angles (Bierhorst, 1977); the spiral phyllotactic pattern observed later likely occurs through another imposed patterning signal. These two examples hint at a position-dependent patterning mechanism which takes place post-apical cell divisions; position-dependent mechanisms tend to be robust and self-organising - just what Thompson might have alluded to above. Evidence for a self-organising and robust patterning mechanism comes from experiments where apical cell ablation does not lead to growth arrest, but instead to a new apical cell establishment and subsequent spiral phyllotaxis about the new centre (Wardlaw, 1949; Cutter, 1965). Work from Wardlaw (1949) and Snow & Snow (1935) explored positional patterning mechanisms which were both physical (tissue tension) and morphogen (the phytohormone auxin) based (reviewed in Philipson, 1990); however, no further modern explorations have been conducted in these species to our knowledge.

In Spermatophytes (seed plants) the meristematic activity in the shoot apex is attributed to an organised group of cells. This niche serves as a reservoir for production of cells which then give rise to the lateral organs (Steeves and Sussex, 1989; Meyerowitz, 1997). Phyllotactic patterning occurs independent from division patterns within the meristematic niche and evidence exists for a position/morphogen-based patterning mechanism: organs emerge due to local softening of tissues at specific positions at the shot apex (Braybrook and Peaucelle, 2013; Milani et al., 2014); local maxima of the phytohormone auxin dictate the position of new organs (Reinhart et al., 2003); auxin maxima are positioned by the cell-cell dynamics of polar auxin transporters whose direction relates to auxin concentration of neighbouring cells (Reinhart et al., 2003); stochastic fluctuations in auxin concentration can lead to coordinated polarisation of auxin transporters and result in a self-organising pattern of organs (Jönsson et al., 2006). Here again, ablation of the meristematic niche leads to re-establishment of a new niche and organised phyllotaxis about the new centre lending weight to a robust self-organising mechanism rooted in the morphogen auxin (Steeves and Sussex, 1989; Reinhardt et al., 2003).

Plants are not the only organisms to display spiral organ arrangement: two genera of parenchymatous multicellular brown algae, in the order Fucales, arrange their organs in spirals: *Sargassum* and *Cystoseira* (Church, 1920). Other members of the order tend to display dichotomous branching (e.g. *Fucus*). The body of parenchymatous brown algae is built through the meristematic activity of an apical cell (Fritsch, 1945; Katsaros, 2000). Most complex brown algal species belong to the order Fucales and have only one apical cell per thallus tip (Yoshida 1983; exceptions detailed in Nizamuddin, 1967; Clayton, 1985). In the Fucales, the apical cell presents as three or four sided in transverse view and divides from these faces (Nizamuddin, 1963; Moss, 1967, 1969; Yoshida, 1983; Klemm and Hallam, 1987; Kaur, 1999). In some cases, the apical cell is thought not to divide but rather stimulate the cells around do so (Moss, 1967). In *Cystophora*, it has been proposed that the division pattern of the apical cell drives the observed branching pattern of the thallus, similar to the theory for moss (Klemm and Hallam, 1987). In *Fucus*, if the apical cell is removed growth of the branch ceases (Moss, 1967) indicating a less robust patterning mechanism than seen in spermatophytes, ferns and lycophytes. As such, the literature indicates that phyllotactic patterns in parenchymatous brown algae may be lineage-dependant.

In *Sargassum muticum*, while a clear apical cell is present (Yoshida, 1983) the shoot apex itself is similar in organisation to that seen in ferns and spermatophytes: a large central area is surrounded by emerging organs in a spiral pattern (Simons, 1906). At the apex a leaf bud is formed when a new apical cell differentiates from the epidermal tissue; subsequently it is proposed that other cells are differentiated into apical cells, between the main apical cell and the leaf apical cell, and these go on to form air bladders and other organs (Oltmann, 1889 via Critchley, 1983). As such the main bud identity observed at the apex is that of a leaf but subsequently branch development occurs, including specification of new lateral branch apices. Since the brown algae have evolved completely independently from plants, it is fascinating to see similar spiral leaf patterns emerging in the shoots of *Sargassum* as are seen in that distant kingdom. Here we explore the apical organisation and spiral phyllotaxis observed in *Sargassum muticum*, and begin to investigate its robustness and underlying mechanisms with comparisons to those proposed in plants.

## Results

### The arrangement of leaf buds in the *Sargassum muticum* meristem follows the golden angle

The *S. muticum* apex has a striking ‘phyllotactic’ pattern, where subsequent branches are spirally organised with respect to each other (Church, 1920). At the apex, these branches begin as leaf buds (Critchley, 1983). In order to characterise the spiral pattern more fully, we performed detailed analysis of *S. muticum* apices collected in the field.

*S. muticum* apices were qualitatively divided into two zones: the apical pit-region, where pro-meristem cells were produced (Fig. 1B, pink), and the peripheral region where new leaf buds formed (Fig. 1B, yellow). In primary lateral apices the meristem size (proxied by presented area of the pit-region, Fig. 1B, pink) was not correlated with stipe length which is representative of apex age (n (individuals) = 7, n (branches) = 22, Fig. S1). Within the peripheral region, the phyllotactic pattern was spiral and presented an average divergence angle (angle between two sequentially-aged buds) of 137.53 ± 2.08° (Fig. 1B, C; n (meristems) = 57). The organisation and phyllotactic pattern observed in the apices of *S. muticum* was highly regular and resembled that seen in complex multicellular plant apices.

**Figure 1.**
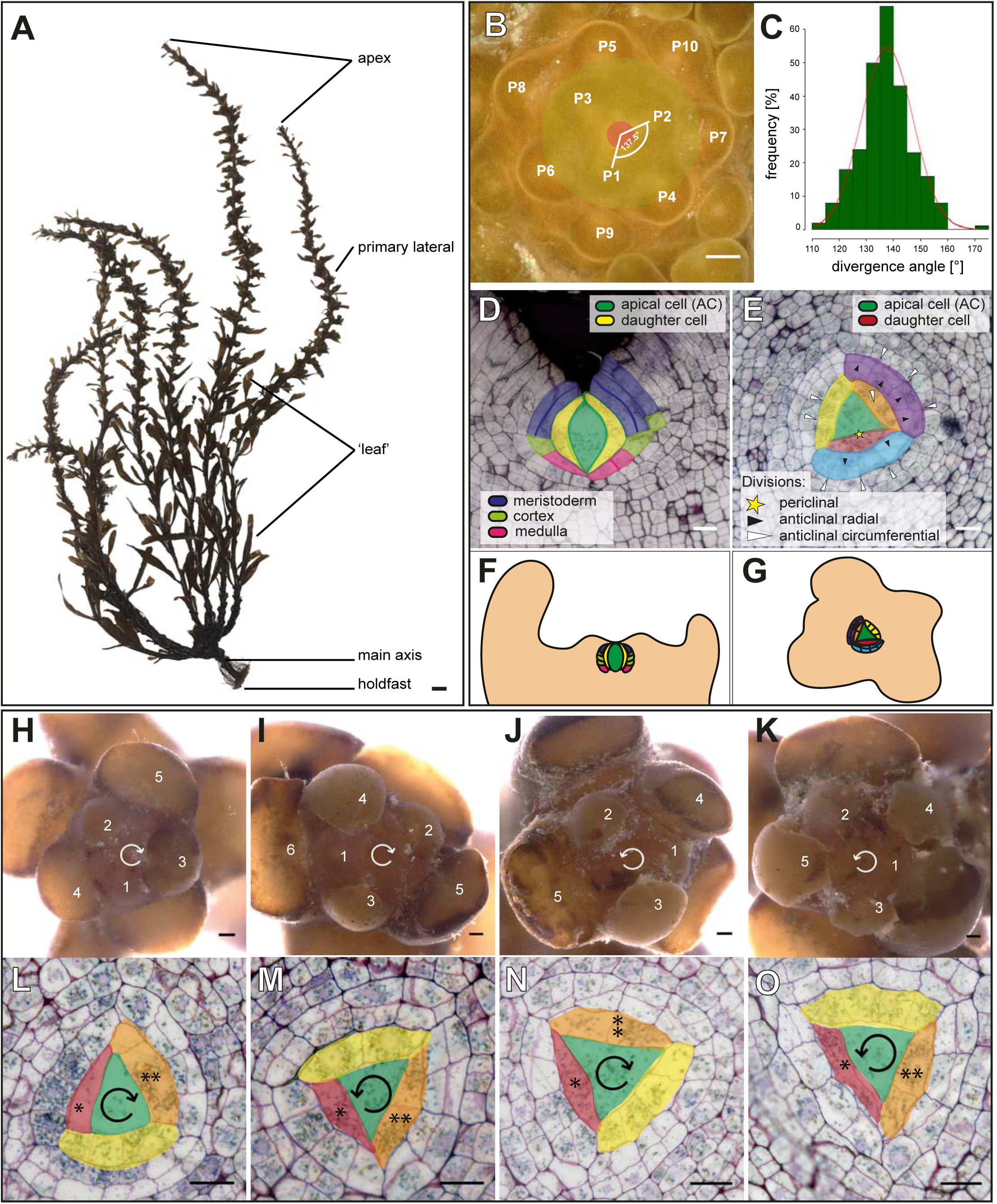
The *Sargassum muticum* apex displays distinct patterns which are independent of each other. (A) The morphology of an adult *S. muticum* alga. (B) Newly forming buds numbered by increasing age (P1 -> P10) with a representative divergence angle illustrated between the two consecutive buds. (C) Divergence angles distribution of measured apices (mean=137.53 ± 2.08°; n=260). (D) Division pattern in a longitudinal section of a *Sargassum* apex; AC divides to give rise to three tissues (meristoderm, cortex, medulla). (E) Apical cell division pattern in a transverse section of a *Sargassum* apex; first periclinal apical cell division (red; yellow star) followed by radial (orange, yellow; white arrowhead) and circumferential (blue; black arrowhead) anticlinal divisions. Schematic representation of the division in the longitudinal direction (F) and the transverse direction (G). (H, I) Clockwise phyllotaxis with a (L) clockwise or (M) counter-clockwise apical cell division orientation. (J, K) Counter-clockwise phyllotaxis with a (N) clockwise or (O) counter-clockwise apical cell division orientation (n=27). *youngest daughter cell, **next-to-youngest daughter cell. Scale bars 1 mm (A), 100 μm (B, H, I, J, K), 20 μm (D, E, L, M, N, O).

### The *Sargassum muticum* apical cell area suggests a highly organised division pattern

As the literature seemed to indicate that brown algae phyllotaxis might be lineage-dependent, we next examined the division patterns of the Sargassum apical cell to see if its pattern exhibited a golden angle, as in moss. The apical cell (AC) of *Sargassum* species has been described as a three-sided lenticel (Yoshida, 1983). In order to investigate the possible patterning of cell divisions in the promeristem, and any connection to the phyllotactic pattern, we examined transverse and longitudinal sections of *S. muticum* apices.

In sections, the apical cell of *S. muticum* presented as bi-convex and lenticular (longitudinally; Fig. 1D) and as three-sided (transversely; Fig. 1E) consistent with other species in the order Fucales (Nizamuddin, 1963; Moss, 1969; Yoshida, 1983; Kaur, 1999). Unlike Fucales apical cells, which are reported to stimulate their neighbours to divide but refrain themselves (Moss, 1967), evidence of apical cell division was observed. In the longitudinal view, divisions appeared to give rise to three tissues – outer layer (meristoderm) and two inner layers (cortex and medulla). The upper cell of the first anticlinal division likely gave rise to the meristodermal and medullar cells whereas the lower cell likely created the medullar and cortex cells (Fig. 1D, F, S2).

In the transverse direction, the first division always appeared to be an asymmetric periclinal division which followed a sequential face division pattern of the apical cell producing promeristem daughters at 119° angles to each other (Fig. 1E, red). Following from this division, the subsequent anticlinal divisions from the initial daughter cell could be followed up to the 6^th^ or 7^th^ ‘division round’ (a division round was defined as a pseudo-time progression that each daughter cell would undergo as it moved away from the apical cell). In the first three rounds of division, the daughter cell then underwent one or more anticlinal radial divisions (Fig. 1E, orange/yellow/blue; white arrowhead); the rounds of radial division never produced more than four cells (Fig. 1E, blue). The further divisions were anticlinal circumferential and created 8 cells in total (Fig. 1E, blue, purple; black arrowhead). These cells then underwent another round of anticlinal radial and circumferential divisions; however, at this point it became difficult to discern lineages in histological sections. The pattern described here was highly conserved although occasionally an anticlinal circumferential division was observed before the 4-cell stage (n=1/30). From these data, it was concluded that the *S. muticum* apical cell divided asymmetrically from sequential faces, producing daughter cells at 120° angles, and that these promeristematic daughter cells further underwent a regimented division pattern.

### The phyllotaxis pattern and the apical cell division pattern are not linked

In *Cystophora*, the apical cell division pattern (bifacial divisions) has been correlated with the apical branching pattern (sympodial branching giving rise to an alternate phyllotactic presentation; Klemm and Hallam, 1987). Our observations in *S. muticum* suggest that the apical cell divides from all three faces to produce promeristem daughter cells at an approximate 120° angle (α=119.01 ± 6.11°, n=74), while the phyllotactic pattern follows at ∼137.5° spiral pattern. In order to examine whether these two patterns in *S. muticum* were linked, we examined the chirality in both the apical and phyllotactic patterns in the same meristems.

The spiral phyllotaxis in *Sargassum muticum* had either a clockwise or a counter-clockwise direction with a ratio of ∼1:1 (58/118 clockwise, 60/118 counter-clockwise). In the clockwise orientation the older buds were located to the left side of the younger bud forming a right-handed spiral (Fig. 1H, I). Likewise, in the counter clockwise orientation, the spiral produced was left-handed (Fig. 1J, K). This 1:1 ratio is observed in plants as well (e.g. Thompson, 1917). With respect to apical cell division patterning, two patterns were observed: moving out from the apical cell, daughter cells were produced to the left or the right yielding both counter- and clockwise patterns in a 1:1 ratio (Fig. 1L, M, N, O; 28/56 clockwise, 28/56 counter-clockwise).

In order to examine if a connection in chirality was observed, individuals were imaged under a light microscope and subsequently sectioned to check the orientation of the apical cell division. In either the counter or clockwise phyllotactic groups, the apical cells presented as ∼1:1 counter- and clockwise (Fig. 1H-O, clockwise phyllotaxis – 8/16 clockwise, 8/16 counter-clockwise apical cell divisions; counter-clockwise phyllotaxis – 7/11 clockwise, 4/11 counter-clockwise apical cell divisions). This data strongly suggests that these two patterning mechanisms are unlinked and may be under separate control. This is highly similar to the patterning mechanisms seen in multicellular plant apices where the phyllotactic pattern is defined in the peripheral zone by the morphogen auxin and the stem cell niche is positioned by another phytohormone, cytokinin (Reinhardt et al., 2003; Chickarmane et al., 2012).

### Ablation of the apical cell leads to formation of a new apical centre indicating pattern self-organisation

Given the observed similarities to multicellular plant meristems, we next examined whether the apical cell and phyllotaxis could re-establish after ablation of the apical cell. In plants, the stem cell region can re-establish in this way pointing to a robust self-organising patterning system (Reinhardt et al., 2003). In *Fucus*, such manipulations led to growth arrest and termination (Moss, 1967).

Apical cell ablations were performed on partially dissected apices using a thin needle aimed at the centre of the pit-area (Fig. 2A, B; white arrowhead). Apices were grown in culture, and re-dissected after a 3-week recovery period before a second imaging. Two scenarios were observed – in 30% (7/23) of the apices the growth of the central zone had stopped or they were dead (6/7 dead, 1/7 no new meristem formed, but the existing buds continued to grow). In 30% of the samples growth continued from what appeared to be a new pit-region (Fig. 2C, D; blue arrowhead, n= 7/23); in these apices, the phyllotactic pattern after recovery exhibited a spiral pattern. In another 20% apices, the meristems seemed to split in two (Fig. 2E; n= 5/23) but again appeared to present spiral patterning. In the remaining 20% of the samples, the results we inconclusive as the imaging methods did not always produce sufficient quality data for pit-area positioning. Culturing itself did not alter the pattern of buds (Fig. 2F).

**Figure 2.**
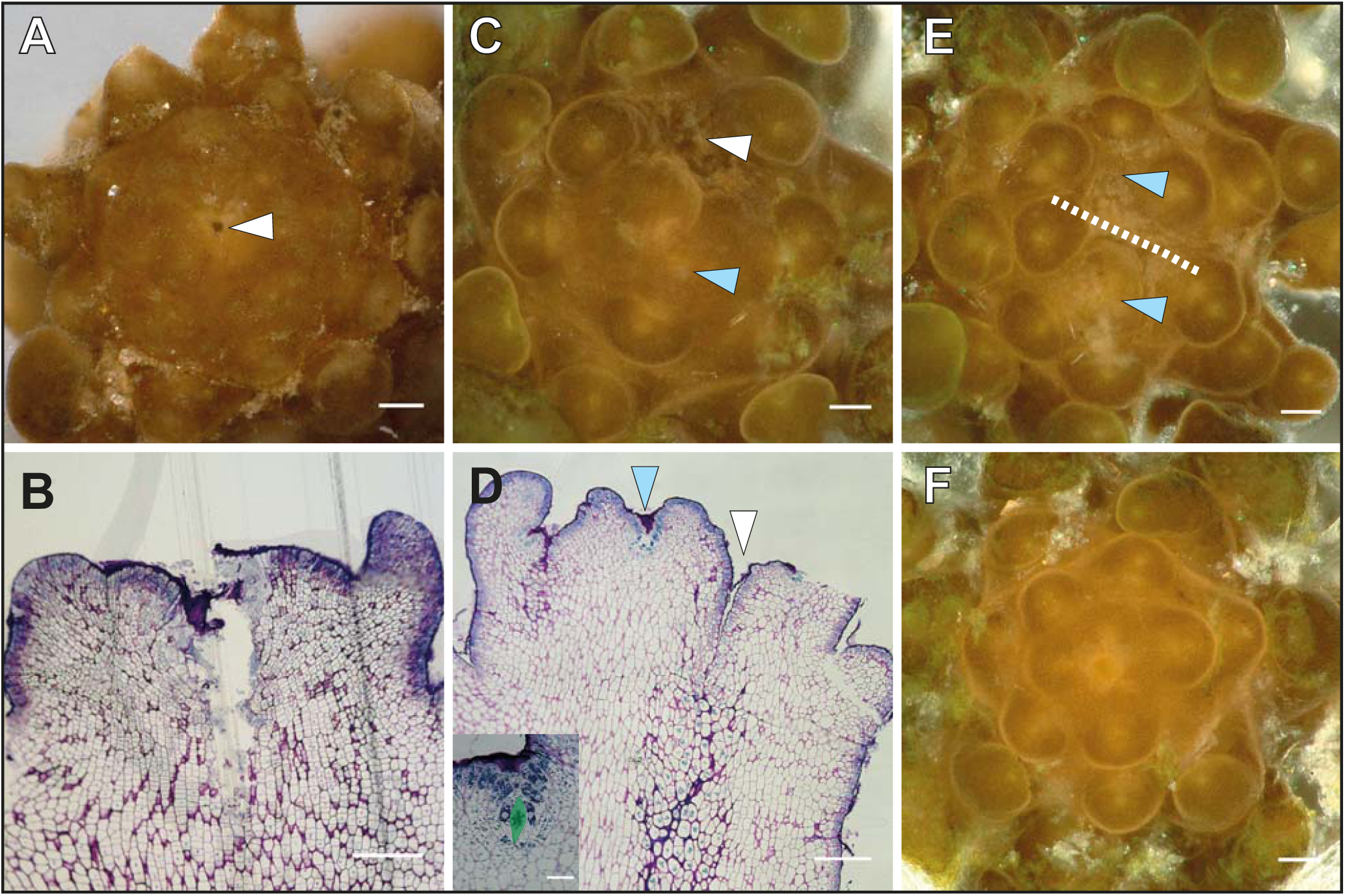
Removal of the apical cell can induce formation of a new central meristem. *Sargassum* meristem stabbed (white arrowhead) in the region of the apical cell in top view (A) and in subsequent longitudinal section (B). (C) Formation of a new central meristem; white arrowhead marks the spot of the stab, blue arrowhead shows the location of the new meristem. (D) Longitudinal section of a newly formed meristem (as in C) showing a new apical cell in the new meristem centre (blue arrowhead) and stabbed area (white arrowhead). Inset: magnified view of the new apical cell. (E) Apex presenting a split meristem; dashed line shows the separation of the two new meristems, centers indicated by blue arrowheads. (F) Control apex, not stabbed. Sample numbers: n=23 stabbed (12 recovered, 7 ceased growth, 4 unclassified). Scale bar 100 μm, 50 μm (B, D), 20 μm (apical cell inset in D).

In the samples where a new pit-area appeared to establish, the wound had moved to the side of the meristem and the new pit-area was roughly centrally positioned (Fig. 2C, D). These data suggest that the meristematic region of the *Sargassum* meristem could re-establish itself indicating a self-organising system similar in nature to that in plant meristems. The data also imply that when a new apical cell is established, the spiral phyllotactic pattern can also re-establish.

### A potential link between auxin and brown algal phyllotaxis is unlikely

In plants, auxin distribution within the peripheral zone dictates the phyllotactic pattern (Reinhardt et al., 2003). Auxin maxima in the peripheral zone lead to cell wall softening and organ outgrowth (Braybrook and Peaucelle, 2013). The brown algae *Fucus vesiculosus* and *Ectocarpus siliquosis* have both exhibited auxin-triggered morphogenetic changes: aberrant embryo rhizoid branching in *Fucus* and filamentous adult branching changes in *Ectocarpus* (Basu et al., 2002; Le Bail et al., 2010). In addition, auxin has been detected using gas chromatography mass spectroscopy and an anti-indole acetic acid (IAA) anti-body in *Ectocarpus* (Le Bail et al., 2010). Since the phyllotaxis in *Sargassum muticum* is spiral and highly resembles the one observed in higher plants and given the potential for auxin response in brown algae, we next examined whether auxin could alter, or be correlated with, the phyllotactic pattern.

We applied auxin externally in the artificial sea water cultivation medium in order to see if phyllotaxis could be altered. In our conditions and experiments, this treatment had no effect on growth or the phyllotactic pattern (50 μM; data not shown). Due to the aqueous nature of the culture system it was not possible to apply auxin locally as has been performed in tomato and *Arabidopsis* (Reinhardt et al., 2000; Braybrook and Peaucelle, 2013). These experiments were therefore inconclusive but lightly suggest that external auxin could not alter phyllotaxis in *Sargassum*, in these conditions.

In order to determine if auxin showed patterned distribution within the apex, we performed immunolocalisations on sectioned apices using the anti-IAA antibody. The anti-IAA signal was strongest in the meristoderm and mucilage external to the meristoderm with accumulation at apical pits and the bases of buds (Fig. 3A, white arrowhead). Internally, there were regions of high signal within the apex in the meristoderm although these did not correlate with bud size or position (Fig. 3A, B). Upon close examination, a large amount of signal originated from the mucilage external to the meristoderm (Fig. 3C). These data suggest that auxin may be accumulating in the meristoderm and mucilage, although its source is undetermined (see Discussion), and there was little correlation with bud position.

**Figure 3.**
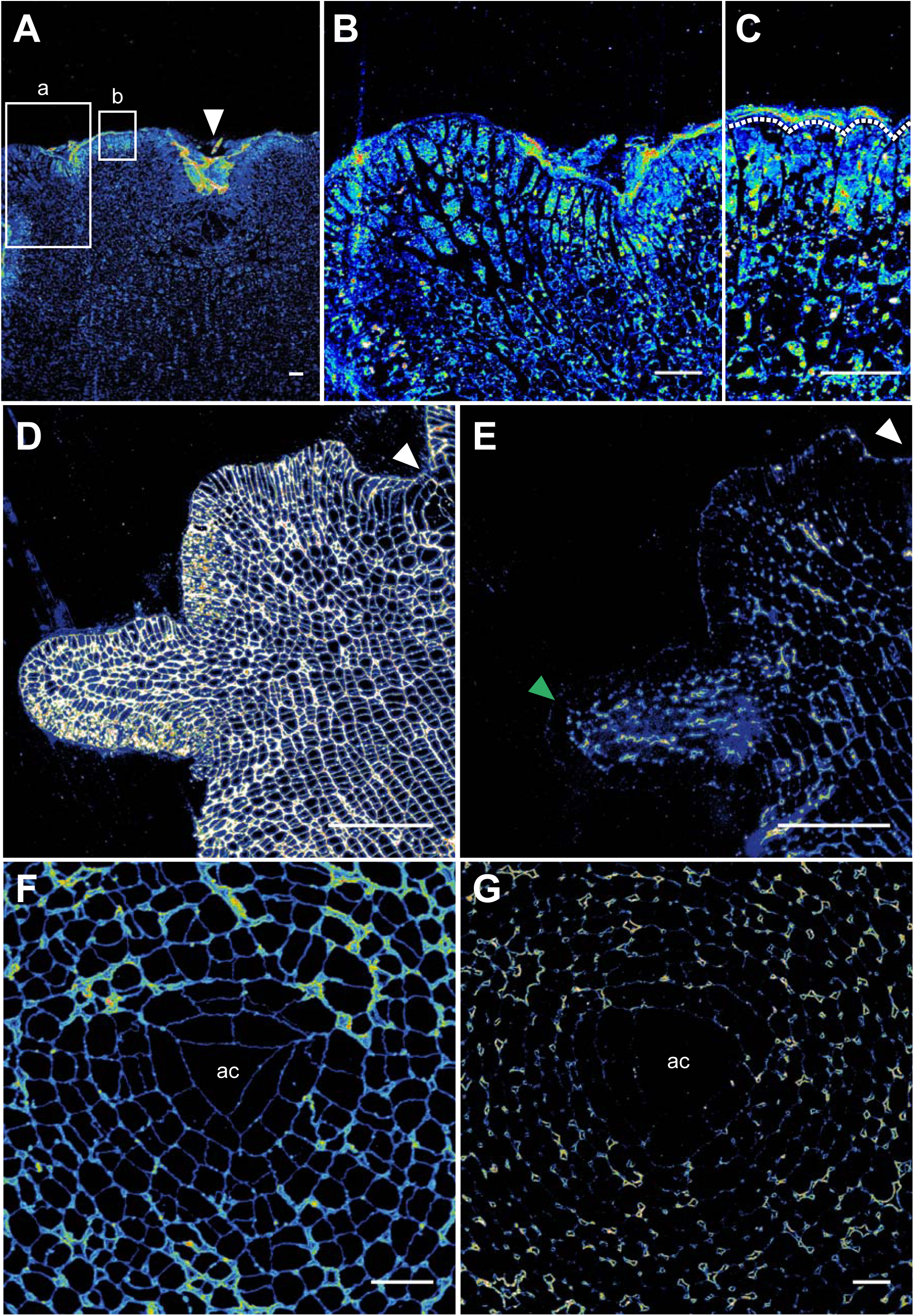
IAA and mannuronate immunolcalisation signals do not appear to correlate with new bud formation in the *S. muticum* apex. (A) Anti-IAA antibody localising to the buds, surface of the meristoderm and the apical pit (white arrowhead). Higher magnification view of the bud (B) and of the meristoderm cells (C). Dashed line in (C) delineates the meristoderm outer cell wall. (D) BAM10 antibody binds to the guluronic acid rich areas in the cell wall with a homogeneous distribution throughout the apex. (E) BAM6 antibody binds to mannuronic acid rich areas in the cell wall, localised at the surface and on cell-cell junctions in the inner tissues and with a slightly higher abundance in a young bud (green arrowhead). (F) BAM10 antibody signal is distributed throughout the apical cell and promeristem region in transverse sections. (G) BAM6 antibody signal is not detected in the apical cell and immediate neighbors. BAM6 localises mainly in the cell junctions around the apical cell. White arrowheads mark the pit-area (with an apical cell). ac= apical cell location. Scale bar 20 μm (A, B, C, F, G), 50 μm (D, E).

### Elongating organs are predicted to have softer walls and the apical cell to have stiffer cell walls

In plants, new organs form in the peripheral zone after auxin maxima lead to wall softening (Braybrook and Peaucelle, 2013). The central zone, containing the meristematic stem cells, exhibits stiffer cell walls than the peripheral zone or new primordia (Milani et al., 2014). In most walled organisms, it is assumed that cell growth is limited by the cell wall and its mechanical properties, in turn linked to its biochemical composition (Peaucelle et al., 2013; Braybrook and Jönsson, 2016). Since *Sargassum* apices were so similar in pattern to the *Arabidopsis* apex, we checked whether they might follow similar mechanical rules. Alginate biochemistry was examined *in muro* using antibodies raised against different biochemical epitopes: the BAM10 antibody preferentially recognises guluronic acid within alginate (Torode et al., 2016), which should be mechanistically rigid through calcium cross-linking (Grant et al., 1973); conversely, the BAM6 antibody preferentially binds to mannuronic acid residues within alginate, which should be mechanically less rigid as they are unable to form calcium cross-links (Torode et al., 2016). Control immunolocalisations may be found in the Supplement (Fig. S3).

In longitudinal sections, BAM10 showed a wide distribution of signal across the apex (Fig. 3D). BAM6 signal was found at the junctions of cells and on the outer periclinal walls of the meristoderm cells (Fig. 3E). There was no obvious pattern of mannuronic/soft alginate associated with young buds, however signal did appear higher in slightly older elongating buds (BAM6; Fig. 3E; green arrowhead). Using these antibodies we were unable to determine if alginate softening could be predicted at the sites of new bud formation; however, it appears that elongating buds have more mannuronic/soft alginate than other areas.

Since the outer wall was thick, and the apical cell covered by a large plug of alginate mucilage (Kaur, 1999; Fig. 1D, dark stained area above the AC), we next looked at the apical cell alginate biochemistry using immunolocalisations on transverse sections. The BAM10 signal was equally distributed across the apical cell, promeristem cells and the surrounding cells (Fig. 3F) in the transverse section. The BAM6 signal was excluded from the apical cell and promeristem cells (Fig. 3G). BAM6 signal was detected in more mature tissues at the junctions between cells (Fig. 3G). These data suggest that the apical cell, young promeristem cells and the peripheral area in the apex have more guluronic acid residues than mannuronic, which may lead to stiffer cell walls.

## Discussion

### Phyllotaxis is a phenomena found in evolutionary distant photosynthetic lineages

Here we report that the spiral phyllotactic pattern in *S. muticum* follows the Golden Angle (∼137.5°) in a pattern almost identical to that found in many multicellular plants. Developmentally, this may not be so surprising: both organisms display indeterminate growth and produce new organs from an apical meristem-like region; both utilise their shoots and branches for light interception; and both utilise branches to produce numerous reproductive structures. However, it is interesting to see a similar pattern when the building blocks of multicellularity are completely independent: while both organisms have cell walls and cell adhesion, the polysaccharides which make up the wall matrix (pectin and alginate) are distinct. These organisms have evolved multicellularity independently (Baldauf, 2008) and therefore may have different communication strategies (morphogen identity): they may be seen as examples of D’Arcy Thompson’s bricklayers whose material is different but results in the same pattern. The data presented here indicate that complex parenchymatous brown algae have position-dependent patterning mechanism which results in spiral phyllotaxis, similar to that observed in plants.

### AC-based patterning does not underlie phyllotactic patterning in *S. muticum*

The apical cell is the centre of the brown algal meristem; its sequential face divisions create a pool of cells which build the body of the adult alga. It has been hypothesised that the division pattern of the algal apical cell directly relates to phyllotactic patterning (Klemm and Hallam, 1987). In the spiral meristem of *S. muticum* this does not appear to be the case. Firstly, the difference in the divergence angles between the two patterns does not support a causative relationship – in apical cell divisions, the angle of the newly produced daughter cell to the previous is 120°, whereas the observed phyllotactic angle centres on the golden angle of 137.5°. In the moss *Atrichum undulatum*, the triangular apical cell exhibits sequential face divisions but these occur at ∼137.5°, and angle which is reflected directly in the phyllotactic angle (Gola and Banasiak, 2016). In *Physcomitrella patens*, spiral apical cell divisions lead to spiral leaf arrangement (Harrison et al., 2009). As this correlation in pattern is not seen in *S. muticum*, apical cell division pattern and phyllotaxis appear to be unrelated. While growth distortion post-apical-cell cannot be discounted, we believe it is unlikely given the highly organised nature of divisions seen in the apex. In fact, the *Sargassum* apex seems more closely aligned with that of the ferns, which also present a three-sided, sequentially-diving, apical cell and robust spiral phyllotaxis (Wardlaw, 1949; Bierhorst, 1977).

A second piece of evidence comes from the observation that both patterns could follow either a left- or a right-handed rotation but the two could be disconnected: a counter-clockwise AC pattern could pair equally with a left- or right-handed phyllotactic spiral. Our experiments suggest that apical cell division pattern and phyllotactic pattern are not correlated. We hypothesise that an apical-cell-independent patterning mechanism exists in *Sargassum*, that position is more instructive than lineage.

### Apical robustness in *S. muticum*

When a stem-cell niche or apical cell are destroyed in plants, there are two outcomes reported in the literature: in ferns the apical growth can cease, or the apex will develop a new stem-cell centre and restart growth (Wardlaw, 1949; Cutter, 1965; Steeves and Sussex, 1989). More recently, in *Solanum lycopersicum* cv *esculentum* a new meristem centre was re-established after laser removal of the original one and the new organs formed in a spiral manner with a ∼137.5° divergence angle (Reinhardt et al., 2003). In brown algae, destruction of the apical cell in leads to the termination of apical growth (Moss, 1967; Clayton and Shankly, 1987); no re-establishment of patterning has been observed to our knowledge.

In our experiments, *S. muticum* apices exhibited both outcomes upon apical-cell destruction: 30% of the apices ceased growing, while another 50% showed continued growth after re-organisation. Our surviving apices fell into two categories: those where a single new apical cell was established or double-meristems where it is likely that two new apical cells were established. Similar to plants, our data showed that the phyllotactic pattern in the *Sargassum* meristem was also re-established (or maintained) upon apical cell destruction. These data indicate that the *S. muticum* apex is capable of re-organisation after apical cell destruction, in a similar way to that seen in plants. This further supports a morphogen-based position-dependent patterning mechanism.

The spiral apices observed in *Sargassum* and *Cystoseira* represent the most complex apices found in the brown algal lineage. This complexity, and its similarity to those of plants, may represent a more robust system when it comes to development: spiral phyllotaxis may allow the algal body a more thorough exploration of space in comparison to the dichotomous thallus found in other Fucales. The ability to re-establish its apical cell and continue patterned growth could also hint at a more robust patterning mechanism in this alga, again as compared to other Fucales.

### Auxin is an unlikely candidate for the phyllotactic morphogen

Hormones are one good candidate for morphogen-based positioning mechanisms in systems with cell walls and fixed position. Auxin’s potential as a morphogen has been established in plants (Reinhardt, 2003) and it has been found to have an effect in the development of Bryophytes (Bennett et al., 2014; Coudert et al., 2015) and brown algae (Basu et al., 2002; Sun et al., 2004; Le Bail et al., 2010). Furthermore, it has been shown in plants that the localisation of auxin at the sites of future primordia changes the cell wall mechanical properties allowing outgrowth to take place (Peaucelle et al., 2011; Braybrook and Peaucelle, 2013). As such, auxin was a prime candidate for morphogen-like behaviour in the *Sargassum* apex.

We were unable to detect an effect of exogenous auxin on phyllotactic patterning in *S. muticum*. In our hands, using a relatively high (but effective in other brown algal cultures; Basu et al., 2002; Le Bail et al., 2010) concentration of auxin we were unable to disturb patterning. It is possible that the auxins used (NAA and 2,4-D) were not effective or that their concentrations require further optimisation. It is also possible that local applications might have been more effective: local applications could alter phyllotaxis in the plants *Lupinus* and *Epilobium* (Snow and Snow, 1937) but when more broadly applied in *Tropaeolum* a lesser effect was seen (Ball, 1944).

In order to gain a more spatial view of possible endogenous auxin, we then switched to IAA-immunlocalisations. Our data suggest that there is no particular localisation of auxin to newly growing buds observed in sections of *S. muticum* apices. The localisation seems to be spread over the whole section with a higher signal localized in the meristoderm cells and mucilage attached to the surface of the alga. We cannot completely rule out a stickiness effect with the mucilage and non-specific antibody reactions; however, the no primary antibody controls were negative (Fig. S3) and the cell wall antibodies did not show such signals (Fig. 3C, D). There are aslso limitations in using an immunolocalisation approach to detect auxin since as it is small and highly dynamic and may be difficult to fix in place; however, using a specific pre-fixation step auxin could be fixed and detected using this approach in plants and brown algae (Avsian-Kretchmer et al. 2002, Le Bail et al. 2010). It is possible that IAA is not the functional auxin in *Sargassum*, although it has been detected by GCMS in the brown alga *Ectocarpus* (Le Bail et al., 2010). The data presented here suggest auxin is an unlikely morphogen for brown algal phyllotaxis, but without a tool such as the molecular reporter constructs in plants we cannot be certain.

The transport of auxin is key to establishing local maxima in plants, while in our algae we did not see evidence of strong maxima. There is no evidence to date that brown algae have homologues of the auxin transporters found in plants (Le Bail et al., 2010; Yue et al., 2014), although the algae may have completely different transport mechanisms. It remains unknown how and if auxins are being transported through the algae. It is always possible that in less complex organisms such as the filamentous *Ectocarpus*, mentioned above, auxin could move by diffusion. Classical phyllotactic patterning mechanisms have relied on reaction-diffusion equations in the past (Bernasconi, 1994; Swinton, 2004), and as such it is possible that auxin diffusion may be instructive in brown algae.

Another question that has been a topic of discussion is whether the auxin detected is being produced by the alga itself rather than being provided by auxin-producing associated bacteria on the surface of their thallus (Evans et al., 1991). Bacteria have been found to mineralize organic substrates giving the algae nutrients and growth factors (Matsuo et al., 2005). They have also been described living in association with brown algae (Hengst et al., 2010; Lachnit et al., 2011), as well as having an effect on their development (Tapia et al., 2016). The marine bacteria *Sulfitobacter* has recently been shown to produce IAA, which is then used by co-incident diatoms to regulate growth (Amin et al., 2015); diatoms are in a sister class to the Phaeophyceae (brown algae). It still remains uncertain how bacteria might be influencing brown algal development, especially when considering about the more complex parenchymous species such as those found in the Fucales. The recent interest in exploring and understanding the algal-bacterial interactions could lead to a better understanding if and how the bacteria might be affecting the development in a more “advanced” brown algal species such as *Sargassum muticum*.

### Cell wall softening and algal bud outgrowth

The cell walls of brown algae are composed mostly out of matrix polysaccharides called alginates and sulphated fucans and a small amount of cellulose (Deniaud-Bouët et al., 2014). It is believed that these are the functional analogues of pectin, hemicelluloses and cellulose in land plants, respectively. In plants, the role of the cell wall and its relationship to auxin in organ formation has been previously explored (Fleming et al., 1997; Reinhardt et al., 1998; Peaucelle et al., 2008, 2011; Braybrook and Peaucelle, 2013), but no information of such is available for the brown algae. Given the recent evidence for a role of pectin in organ emergence, and the predominant nature of alginate in the algal cell wall it seems plausible that alginate may be involved in bud formation.

Alginate is formed of two residues, mannuronic and guluronic acid, the latter being able to cross-link with Ca^2+^ ions (Grant et al., 1973), similarly to pectin in plants, and thus change its mechanical properties (Mancini et al., 1999). New alginate is added to the wall in the softer, mannuronic acid, form and may then be selectively epimerised into the guluronic acid form which can calcium cross-link (Grant et al., 1973). New techniques have been recently developed to look into the brown algal cell wall biochemistry by using immunolocalisations targeted towards specific epitopes in alginate chains (Torode et al., 2016). We have used two antibodies which bind to two different alginate epitopes, one rich in guluronic acid (BAM10) and the other rich in mannuronic acid residues (BAM6). Our data suggests that the guluronic acid rich areas are more abundant throughout the apex of *S. muticum*, but there is no clear distinction between growing and non-growing parts. The mannuronic acid (BAM6) signal tended to be higher in rapidly growing older buds but did not obviously mark young buds. These observations differ from the ones seen in *A. thaliana* meristems where pectin biochemical changes did mark new organ sites (Peaucelle et al., 2011). In slightly older, elongating buds, a stronger signal from BAM6 was detected, indicating a possible role for alginate biochemistry in elongation but not initiation of buds.

When looking into the apical cell and the cells around it in transverse sections, we observed that the guluronic acid signal (BAM10) was high whereas that for mannuronic acid (BAM6) was barely detected. Based on these data, we could suggest that the area around the apical cell and the apical cell itself have stiffer walls. This is similar to the situation observed in plants, where the stem cells have been shown to be stiffer than the surrounding peripheral cells (Milani et al., 2014). It may be interesting to explore whether this increased stiffness regulates cell division rates, in both plants and algae.

Taken together, it seems that the apical cell in *Sargassum* may behave mechanically similar to the stem-cell niche in spermatophyte plants and that softer alginate may be present in rapidly elongating new buds. However, it does not appear that new bud positions display biochemical markers of softer alginate and as such it is possible that alginate biochemistry is not related to bud positioning in *Sargassum*, only in outgrowth.

### Possible mechanisms of phyllotaxis in *S. muticum*

In the *Sargassum* apex, a new leaf will develop its own apical cell, and further cells along the meristoderm between this and the primary apical cell will follow suit, each giving rise to another organ on each branch (Oltmann 1889 via Critchley 1983). All meristodermal cells have a meristematic ability which could indicate that any cell from this cell layer could “switch on” and become an apical cell and start producing its own bud (Moss, 1967). This is not dissimilar to the specification seen in ferns for the production of new leaves from the epidermis (Bierhorst, 1977; Mueller, 1982). Based on the robust, self-organising, nature of the *Sargassum* apex and the lack of correlation between the apical cell division pattern and that of new buds, it seems likely that a positional mechanism is in place for phyllotactic patterning in *Sargassum*. It is then plausible that an unknown morphogen instructs the conversion of a meristodermatic cell into a new apical one, in a positioned manner. The secondary specification of further apical cells between the leaf and main apical cell also hints at a position-dependent specification of meristodermal conversion into apical cells.

It has been observed that the apical cell can divide to potentially create a daughter apical cell which then continues to create a new branch (Kaur, 1999). In our experiments, we never observed an equal division of the apical cell which could explain the previously described situation. This is similar to the case in *Cystophora* where division to produce a second apical cell was rarely observed (Klemm and Hallam, 1987). If we assume this hypothesis might be true and that we simply missed such special divisions, based on the observed patterns of longitudinal division it seems unlikely that this could produce a golden-angled spiral and would more likely produced a 120° spiral.

The absence of cell wall biochemical marks associated with alginate softening (mannuronic vs. guluronic acid) correlated to new bud positions indicates that the physical events of initial bud outgrowth may be different than that in plants. This does not rule out a physically-based positioning system for the brown algal apex; physical buckling may give rise to phyllotactic patterns. While the mechanical properties of the meristoderm remain homogeneous, if the underlying cortex tissue is growing at a differential rate to the meistoderm physical buckling may result through compression of the outer tissue (reviewed in Dumais, 2007). This possibility is worthy of further investigation.

### Future directions

The similarity between the *Sargassum* apex and that of complex multicellular plant meristems is striking: the presence of a golden-angled phyllotactic spiral; the robust reorganisation after meristem ablation; the presence of equal clockwise and counter-clockwise patterns; the apparent independence of phyllotactic patterning to meristem divisions. However, there are many obvious differences as well: we do not currently have strong evidence for auxin as a patterning morphogen; we also cannot detect softening of the algal cell walls coincident with new bud outgrowth.

While the experiments presented here make a case for the *Sargassum* apex as being more plant-like in its patterning and organisation principals, we have many new questions. Is there a re-specification of a meristodermal cell into a new apical cell, and how is this regulated? If auxin is an unlikely morphogen, what might be the identity of the algal functional analogue (if it exists at all)? If auxin can in fact be instructive, is it produced by the algae or by associated bacteria (implying a more communal evolution of patterning in the brown algae)?

The answers to many of these questions undoubtedly require advances in molecular techniques and genetics within the brown algae. These techniques are beginning to be developed in *Ectocarpus* and hopefully can be translated into other interesting algae (Le Bail et al., 2011). Another hurdle is the inability to culture many brown algae for their full life cycle *in vitro*, thus limiting when questions might be asked (seasonally). Currently methods exist for *Ectocarpus* and *Dictyota* (Le Bail et al., 2011; Bogaert et al., 2016). In addition, it remains unclear how axenic growth conditions can become in this cultures while still supporting growth and development.

## Materials and Methods

### Sample collection and *in vitro* culture

The samples were collected in Rottingdean (East Sussex, United Kingdom) between November 2015 and February 2017. After collection, they were transported in seawater to the laboratory and kept at 4°C. Processing was done shortly after returning back to the laboratory – the specimens were dissected using fine tweezers and used for further experiments. The medium used in the experiments was filter sterilised artificial seawater (ASW, Tropic Marin Sea Salt; Tropic Marin, Germany).

### Imaging of the apices for divergence angle measurements

*Sargassum* apices were dissected using fine tweezers by removing all the leaves from their base, until the central region of the apex was clearly visible. The apices were then cut to a 1 cm length and anchored by insertion into Petri dishes containing 1% agarose melted in ASW and flooded with ASW to cover. Images were taken using a VHX 5000 microscope (Keyence (UK) Ltd, UK). Measurements for the divergence angle were done using the VHX 5000 Keyence software; centres of each organ were used in reference to the centre of the apex.

### Histology

The apices were fixed in a fixative containing 2.5% glutaraldehyde and 2% formaldehyde in artificial seawater. They were then dehydrated through 10% ethanol steps and embedded in resin (LR White resin, Agar Scientific Ltd, UK). Samples were then cut in 1 μm slices using a Leica EM UC7 ultramicrotome (Leica Microsystems, Germany). Sections were placed onto Superfrost Ultra Plus slides (Thermo Scientific, USA) and left to dry at room temperature. Sections were then stained with 0.05% Toluidine blue O solution for 5 minutes, washed, covered with a cover slip and imaged under the Zeiss Axio Imager M2 (Zeiss, Germany).

### Apex ablation

*Sargassum* apices were dissected and handled as described above. They were then precisely stabbed using a fine needle in the middle of the meristem, where the apical cell is located. The images of the stabbed meristems were taken using a VHX 5000 microscope (Keyence (UK) Ltd, UK). The samples were kept in culture under 16°C, 12:12 hour day night cycle, 60 μmol m^-2^ s^-1^. After 20 days, they were dissected again to remove the newly grown leaves and imaged again. They were fixed, dehydrated, embedded in resin as previously described. The embedded apices were then cut into serial 1 μm sections (every 5 μm throughout the meristem), stained with TBO as above and imaged under Zeiss Axio Imager M2 (Zeiss, Germany).

### Alginate immunolocalisation

The apices were fixed, dehydrated and embedded in resin as described above. The sections were placed on Vectabond coated multitest 8-well slides, 2 sections per well (Vector Laboratories, USA). They were then incubated in a blocking solution of 5% milk for 2 hours. The sections were rinsed with phosphate buffered saline (PBS; 2.7 mM KCl, 6.1 mM Na_2_HPO_4_, and 3.5 mM KH_2_PO_4_) and incubated in the 60 μl of 1/5 (in 5% milk) monoclonal primary antibody for 1.5 hours. After the incubation, the slides were washed with PBS 3 times for 5 minutes each, followed by the incubation in the 60 μl of 1/100 (in 5% milk) IgG-FITC secondary antibody (F1763, Sigma-Aldrich). The sections were then washed 5 times for 5 minutes in PBS, mounted in Citifluor (Agar Scientific, UK), covered with a coverslip, sealed and imaged under a Leica SP8 confocal microscope (Leica Microsystems, Germany). The antibodies used were gifts from Prof. Paul Knox (University of Leeds).

### Auxin immunolocalisation

The protocol was adapted from Le Bail et al. (2010). The *Sargassum* and *Arabidopsis* apices were dissected and prefixed in 3% of 1-ethyl-3-(3-dimethylaminopropyl)-carbodiimide (EDAC, Sigma-Aldrich, USA) followed by an overnight fixation in FAA ((47.5% ethanol, 5%acetic acid, and 10% formaldehyde in ASW). Samples were then dehydrated and embedded in resin as described above. 1μm sections were cut using the Leica ultramicrotome and placed on Vectabond coated slides. The slides were placed into PBS for 5 minutes and then incubated in a blocking solution (0.1% [v/v] Tween 20, 1.5% [w/v] Glycine, and 5% [w/v] bovine serum albumin (BSA) in dH_2_O) for 45 minutes. The sections were rinsed in a salt rinse solution for 5 minutes, a quick wash with 0.8% (w/v) BSA in PBS and incubated in 60 μl of 1:100 monoclonal anti-IAA antibody (Sigma Aldrich, USA) overnight at 4°C. The slides were vigorously washed three times for 10 minutes with a high salt rinse solution (2.9% [w/v] NaCl, 0.1% [v/v] Tween 20, and 0.1% [w/v] BSA in dH_2_O) and then washed for an additional 10 minutes in a salt rinse solution and a rinse with 0.8% (w/v) BSA and then in PBS. 60 μl of 1:100 (v/v) dilution of the 1mgmL^-1^ goat anti-mouse IgG antibody Alexa Fluor^®^ 488 (Invitrogen, USA) was added to each well and incubated for 4 h at room temperature. The slides were washed 5 times for 10 minutes in the salt rinse solution followed by a brief was in PBS, mounted in Citifluor (Agar Scientific, UK), covered, sealed and imaged under a Leica SP8 confocal microscope (Leica Microsystems, Germany).

## Acknowledgements

We thank Dr. Thomas Torode for help with immunolocalisations, Dr. Paul Knox for the BAM10 and BAM6 antibodies, and the Braybrook Group for helpful feedback on the project. We thank our colleagues in the Phycomorph Network for their support and collaboration (COST Action FA1406; http://www.phycomorph.org/).

## Data availability

All raw data and images are available from the corresponding author upon request.

---

**Supplementary Figure 1.**
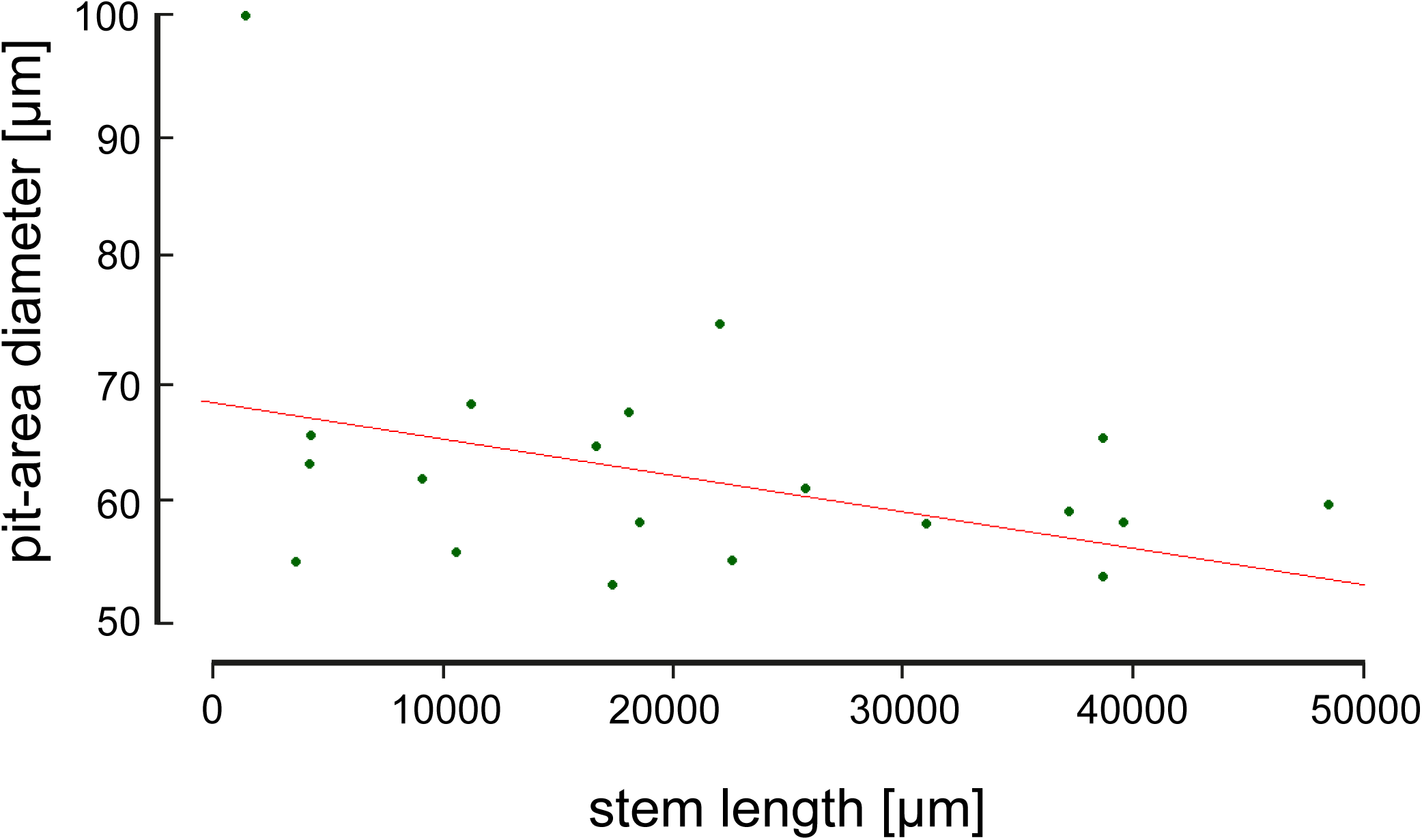
The age of the stipe and the meristem area. Scatter plot showing a lack of correlation between the length (proxy for age) of an individual stipe with the diameter of its pit-area (proxy for meristem size; apical cell and the promeristem cells around it) (n=22, p-value=0.07, r=-0.39; two-sample t-test).

**Supplementary Figure 2.**
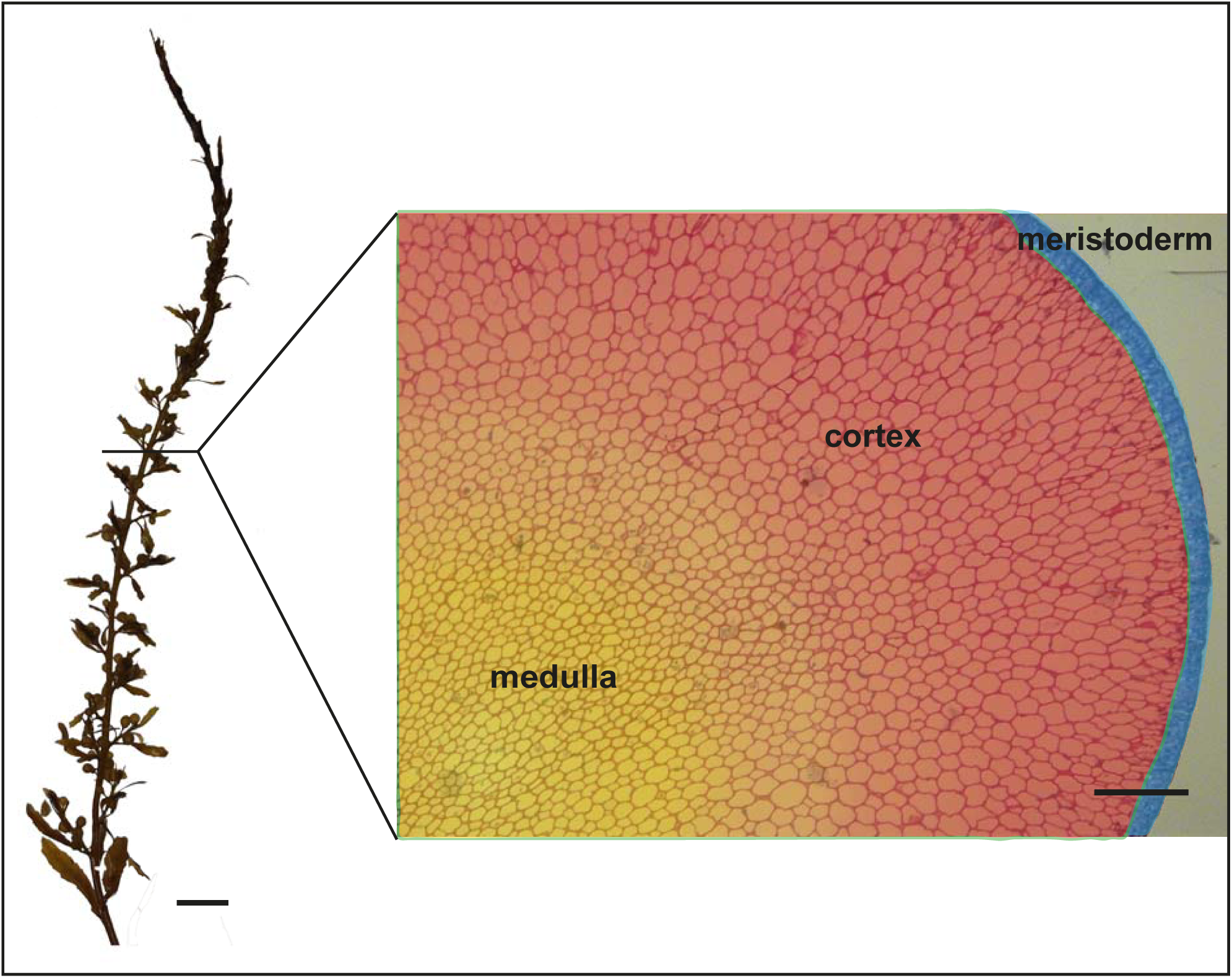
Types of tissues found in *S.muticum* as illustrated on a stipe section. Outer layer (meristoderm), middle layer (cortex) and inner layer (medulla). Scale bar 150 μm, 1 mm (whole algal body).

**Supplementary Figure 3.**
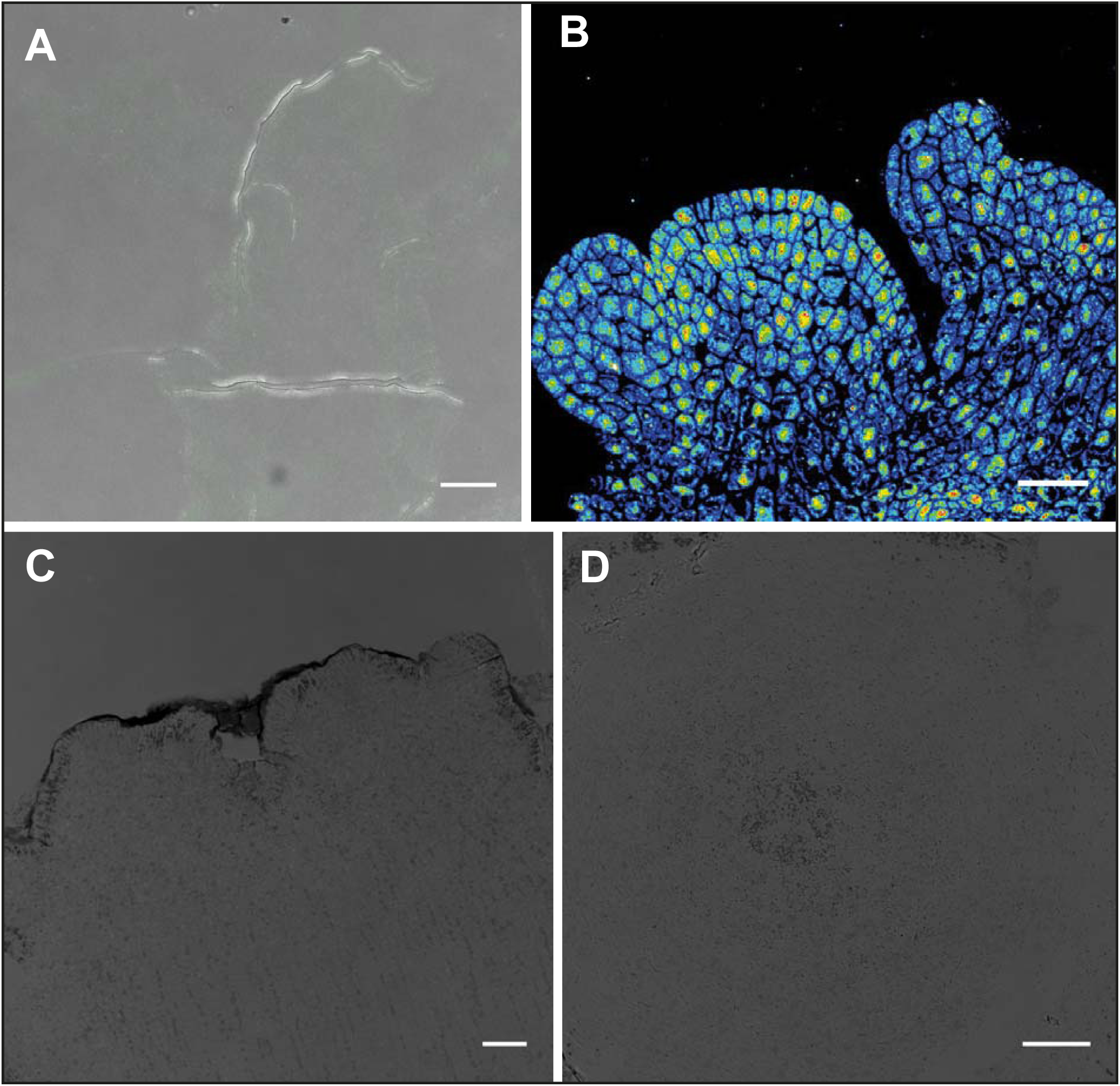
Control confocal images for the auxin and alginate immunolocalisations. *Arabidopsis thaliana* longitudinal section with no primary antibody control (A) and anti-IAA (B). (C) No primary antibody controls of *S. muticum* apex sections for alginate immunolocalisation (C) longitudinal section and (D) transverse. All controls merged with a brightfield image for visualisation. Scale bar 20 μm.

## References

Amin, S. A., Hmelo, L. R., Van Tol, H. M., Durham, B. P., Carlson, L. T., Heal, K. R., Morales, R. L., Berthiaume, C. T., Parker, M. S., Djunaedi, B., et al. (2015). Interaction and signalling between a cosmopolitan phytoplankton and associated bacteria. Nature 522, 98–101.

Avsian-Kretchmer, O., Cheng, J. C., Chen, L., Moctezuma, E. and R, S.Z. (2002). Indole Acetic Acid Distribution Coincides with Vascular Differentiation Pattern during Arabidopsis Leaf Ontogeny. Plant Physiol. 130, 199–209.

Baldauf, S. L. (2008). An overview of the phylogeny and diversity of eukaryotes. J. Syst. Evol. 46, 263–273.

Ball, E. (1944). The Effects of Synthetic Growth Substances on the Shoot Apex of Tropaeolum majus L. Am. J. Bot. 31, 316–327.

Basu, S., Sun, H., Brian, L., Quatrano, R. L. and Muday, G. K. (2002). Early Embryo Development in Fucus distichus Is Auxin Sensitive. Plant Physiol. 130, 292–302.

Bennett, T. A., Liu, M. M., Aoyama, T., Bierfreund, N. M., Braun, M., Coudert, Y., Dennis, R. J., O’Connor, D., Wang, X. Y., White, C. D., et al. (2014). Plasma Membrane-Targeted PIN Proteins Drive Shoot Development in a Moss. Curr. Biol. 24, 2776–2785.

Bernasconi, G. P. (1994). Reaction-diffusion model for phyllotaxis. Phys. D 70, 90–99.

Bierhorst, D. W. (1977). On the Stem Apex, Leaf Initiation and Early Leaf Ontogeny in Filicalean Ferns. Am. J. Bot. 64, 125–152.

Bogaert, K., Beeckman, T. and De Clerck, O. (2016). Abiotic regulation of growth and fertility in the sporophyte of Dictyota dichotoma (Hudson) J.V. Lamouroux (Dictyotales, Phaeophyceae). J. Appl. Phycol.

Braybrook, S. A. and Jönsson, H. (2016). Shifting foundations: The mechanical cell wall and development. Curr. Opin. Plant Biol. 29, 115–120.

Braybrook, S. A. and Peaucelle, A. (2013). Mechano-Chemical Aspects of Organ Formation in Arabidopsis thaliana: The Relationship between Auxin and Pectin. PLoS One 8, e57813.

Chickarmane, V. S., Gordon, S. P., Tarr, P. T., Heisler, M. G. and Meyerowitz, E. M. (2012). Cytokinin signaling as a positional cue for patterning the apical–basal axis of the growing Arabidopsis shoot meristem. Proc. Natl. Acad. Sci. 109, 4002– 4007.

Church, A. H. (1920). On the interpretation of phenomena of phyllotaxis. London: Humphrey Milford and Oxford University Press.

Clayton, M. N. and Shankly, C. M. (1987). The apical meristem of Splachnidium rugosum (Phaeophyta). J. Phycol. 23, 296–307.

Clayton, M. N. (1985). A critical investigation of the vegetative anatomy, growth and taxonomic affinities of Adenocystis, Scytothamnus, and Splachnidium (Phaeophyta). Br. Phycol. J. 20, 285–296.

Coudert, Y., Palubicki, W., Ljung, K., Novak, O., Leyser, O. and Harrison, J. C. (2015). Three ancient hormonal cues co-ordinate shoot branching in a moss. Elife 4, e06808.

Critchley, A. T. (1983). Sargassum muticum: a morphological description of european material. J. Mar. Biol. Assoc. United Kingdom 63, 813–824.

Cutter, E. G. (1965). Recent Experimental Studies of the Shoot Apex and Shoot Morphogenesis. Bot. Rev. 31, 7–113.

Deniaud-Bouët, E., Kervarec, N., Michel, G., Tonon, T., Kloareg, B. and Hervé, C. (2014). Chemical and enzymatic fractionation of cell walls from Fucales: Insights into the structure of the extracellular matrix of brown algae. Ann. Bot. 114, 1203– 1216.

Dumais, J. (2007). Can mechanics control pattern formation in plants? Curr. Op. Plant Biol. 10, 58–62.

Evans, L. V and Trewavas, A. J. (1991). Is algal development controlled by plant growth substances? J. Phycol. 27, 322–326.

Fleming, A. J. (1997). Induction of Leaf Primordia by the Cell Wall Protein Expansin. Science (80-.). 276, 1415–1418.

Fritsch, F. E. (1945). Observations on the anatomical structure of the Fucales. New Phytol. 44, 1–16.

Gola, E. M. and Banasiak, A. (2016). Diversity of phyllotaxis in land plants in reference to the shoot apical meristem structure. Acta Soc. Bot. Pol. 85, 3529.

Golub, S. J. and Wetmore, R. H. (1948). Studies of Development in the Vegetative Shoot of Equisetum arvense L. I. The Shoot Apex. Am. J. Bot. 35, 755–767.

Grant, G. T., Mon, E. R., Rees, D. A., Smith, P. J. C. and Thom, D. (1973). Biological interactions between polysaccharides and divalent cations: the egg-box model. FEBS Lett. 32, 195–198.

Harrison, C. J., Roeder, A. H. K., Meyerowitz, E. M. and Langdale, J. A. (2009). Local Cues and Asymmetric Cell Divisions Underpin Body Plan Transitions in the Moss Physcomitrella patens. Curr. Biol. 19, 461–471.

Hengst, M. B., Andrade, S., González, B. and Correa, J. A. (2010). Changes in Epiphytic Bacterial Communities of Intertidal Seaweeds Modulated by Host, Temporality, and Copper Enrichment. Microb. Ecol. 60, 282–290.

Jönsson, H., Heisler, M. G., Shapiro, B. E., Meyerowitz, E. M. and Mjolsness, E. (2006). An auxin-driven polarized transport model for phyllotaxis. Proc. Natl. Acad. Sci. 103, 1633–1638.

Katsaros, C. I. (1995). Apical cells of brown algae with particular reference to Sphacelariales, Dictyotales and Fucales. Phycol. Res. 43, 43–59.

Kaur, I. (1999). Apical meristem of Sargassum vulgare C. Agardh (Phaeophyta,Fucales). Algae 14, 37–42.

Klemm, M. F. and Hallam, N. D. (1987). Branching pattern anf growth in Cystophora (Fucales, Phaeophyta). Phycologia 26, 252–261.

Korn, R. W. (1993). Apical cells as meristems. Acta Biotheor. 41, 175–189.

Lachnit, T., Meske, D., Wahl, M., Harder, T. and Schmitz (2011). Epibacterial community patterns on marine macroalgae are host-specific but temporally variable. Environ. Microbiol. 13, 655–665.

Le Bail, A., Billoud, B., Kowalczyk, N., Kowalczyk, M., Gicquel, M., Le Panse, S., Stewart, S., Scornet, D., Cock, J. M., Ljung, K., et al. (2010). Auxin metabolism and function in the multicellular brown alga Ectocarpus siliculosus. Plant Physiol. 153, 128–44.

Le Bail, A., Billoud, B., Le Panse, S., Chenivesse, S. and Charrier, B. (2011). ETOILE regulates developmental patterning in the filamentous brown alga Ectocarpus siliculosus. Plant Cell 23, 1666–1678.

Mancini, M., Moresi, M. and Rancini, R. (1999). Mechanical properties of alginate gels: empirical characterisation. J. Food Eng. 39, 369–378.

Matsuo, Y., Imagawa, H., Nishizawa, M. and Shizuri, Y. (2005). Isolation of an Algal Morphogenesis Inducer from a Marine Bacterium. Science (80-.). 307, 1598.

Meyerowitz, E. M. (1997). Genetic Control Review of Cell Division Patterns in Developing Plants. Cell 88, 299–308.

Milani, P., Mirabet, V., Cellier, C., Rozier, F., Hamant, O., Das, P. and Boudaoud, A. (2014). Matching Patterns of Gene Expression to Mechanical Stiffness at Cell Resolution through Quantitative Tandem Epifluorescence and Nanoindentation. Plant Physiol. 165, 1399–1408.

Moss, B. (1969). Apical Meristems and Growth Control in Himanthalia Elongata (S. F.Gray). New Phytol. 68, 387–397.

Moss, B. (1967). The apical meristem of Fucus. New Phytol. 66, 67–74.

Mueller, R. J. (1982). Shoot Morphology of the Climbing Fern Lygodium (Schizaeaceae): General Organography, Leaf Initiation, and Branching. Bot. Gaz. 143, 319–330.

Nägeli, C. (1845). Wachtumsgeschiehte yon Delesseria Hypoglossum. Zeitschrit. Wiss. Bot 2, 121–137.

Nägeli, C. (1845). Wachtumsgesehichte der Laubund Lebermoose. Zeitschrit. Wiss. Bot 2, 138–210.

Nizamuddin, M. (1967). Morphology and Anatomy of Phyllospora, Scytothalia and Seirococcus (Fucales). Bot. Mar. 81–105.

Oltmann, S. F. (1889). Beitrage zur Kenntniss der Fucaceen. Bild. Bot. 14.

Peaucelle, A., Braybrook, S. A., Le Guillou, L., Bron, E., Kuhlemeier, C. and Höfte, H. (2011). Report Pectin-Induced Changes in Cell Wall Mechanics Underlie Organ Initiation in Arabidopsis. Curr. Biol. 21, 1720–1726.

Peaucelle, A., Louvet, R., Johansen, J. N., Höfte, H., Laufs, P., Pelloux, J. and Mouille, G. (2008). Arabidopsis Phyllotaxis Is Controlled by the MethylEsterification Status of Cell-Wall Pectins. Curr. Biol. 18, 1943–1948.

Philipson, W. R. (1990). The significance of apical meristems in the phylogeny of land plants. Plant Syst. Evol. 173, 17–38.

Reinhardt, D., Frenz, M., Mandel, T. and Kuhlmeier, C. (2003). Microsurgical and laser ablation analysis of interactions between the zones and layers of the tomato shoot apical meristem. Development 130, 4073–4083.

Reinhardt, D., Mandel, T. and Kuhlmeier, C. (2000). Auxin Regulates the Initiation and Radial Position of Plant Lateral Organs. Plant Cell 12, 507–518.

Reinhardt, D., Pesce, E. R., Stieger, P., Mandel, T., Baltensperger, K., Bennett, M., Traas, J., Friml, J. and Kuhlemeier, C. (2003). Regulation of phyllotaxis by polar auxin transport. Nature 426, 255–260.

Reinhardt, D., Wittwer, F., Mandel, T. and Kuhlemeier, C. (1998). Localized upregulation of a new expansin gene predicts the site of leaf formation in the tomato meristem. Plant Cell 10, 1427–1437.

Renzaglia, K. S., Dui, R. J., Nickrent, D. L. and Garbary, D. J. (2000). Vegetative and reproductive innovations of early land plants: implications for a unifed phylogeny. Philos. Trans. R. Soc. B, 769–793.

Scheres, B. (2001). Plant cell identity. The role of position and lineage. Plant Physiol. 125, 112–114.

Schüepp, O. (1926). Meristeme. In Handbuch der Pflanzenanatomie, p. Berlin: Gebrüder Borntraeger.

Simons, E. B. (1906). A morphological study of Sargassum filipendula. Bot. Gazzette 41, 161–182.

Snow, M. and Snow, R. (1935). Experiments on Phyllotaxis. Part III. Diagonal Splits through Decussate Apices. Philos. Trans. R. Soc. B Biol. Sci. 225, 63–94.

Steeves, T. and Sussex, I. (1989). Patterns in Plant Development. UK: Cambridge University Press.

Sun, H., Basu, S., Brady, S. R., Luciano, R. L. and Muday, G. K. (2004). Interactions between auxin transport and the actin cytoskeleton in developmental polarity of Fucus distichus embryos in response to light and gravity. Plant Physiol. 135, 266–278.

Swinton, J. (2004). Watching the Daisies Grow: Turing and Fibonacci Phyllotaxis. In Alan Turing: Life and Legacy of a Great Thinker (ed. Teuscher, C.), pp. 477–498. Berlin, Heidelberg: Springer Berlin Heidelberg.

Tapia, J. E., González, B., Goulitquer, S., Potin, P. and Correa, J. A. (2016). Microbiota influences morphology and reproduction of the brown alga *Ectocarpus* sp. Front. Microbiol. 7.

Thompson, D. W. (1917). On Growth and Form. UK: Cambridge University Press.

Torode, T. A., Siméon, A., Marcus, S. E., Jam, M., Le Moigne, M. A., Duffieux, D., Knox, J. P. and Hervé, C. (2016). Dynamics of cell wall assembly during early embryogenesis in the brown alga Fucus. J. Exp. Bot.

Wardlaw, C. W. (1949). Phyllotaxis and organogenesis in ferns. Nature 167–169.

Yoshida, T., Majima, T. and Marui, M. (1983). Apical organization of some genera of Fucales (Phaeophyta) from Japan. J. Fac. Sci. Hokkaido Univ. 13, 49–56.

Yue, J., Hu, X. and Huang, J. (2014). Origin of plant auxin biosynthesis. Trends Plant Sci. 19, 764–770.

